# Sensory target detection at local and global timescales reveals a hierarchy of supramodal dynamics in the human cortex

**DOI:** 10.1101/2022.02.02.478851

**Authors:** Maria Niedernhuber, Federico Raimondo, Jacobo D. Sitt, Tristan A. Bekinschtein

**Author notes:** Author Note: Maria Niedernhuber and Federico Raimondo share first authorship. Tristan Bekinschtein and Jacobo D. Sitt share last authorship. Preregistration: https://osf.io/3mqvy/. The authors declare no competing financial interests. The authors made the following contributions. Maria Niedernhuber: Conceptualization, Data collection, Data analysis, Writing - Original Draft Preparation, Writing - Review & Editing; Federico Raimondo: Conceptualization, Data collection, Data analysis, Writing - Review & Editing; Jacobo D. Sitt: Conceptualization, Writing - Review & Editing; Tristan A. Bekinschtein: Conceptualization, Writing - Review & Editing. Correspondence concerning this article should be addressed to Maria Niedernhuber, Craik-Marshall Building, 20 Downing Pl, Cambridge CB2 1QW, United Kingdom.

## Abstract

To ensure survival in a dynamic environment, the human neocortex monitors input streams forwarded from different sensory organs for important sensory events. Which principles govern whether different senses share common or modality-specific networks for sensory target detection? We examined whether complex targets evoke sustained supramodal activity while simple targets rely on modality-specific networks with short-lived supramodal contributions. In a series of hierarchical multisensory target detection studies (n=77, of either sex) using Electroencephalography, we applied a temporal cross-decoding approach to dissociate supramodal and modality-specific cortical dynamics elicited by rule-based global and feature-based local sensory deviations within and between the visual, somatosensory and auditory modality. Our data show that each sense implements a cortical hierarchy which orchestrates supramodal target detection responses operating on local and global timescales at successive processing stages. Across different sensory modalities, simple feature-based sensory deviations presented in temporal vicinity to a monotonous input stream triggered an MMN-like local negativity which decayed quickly and early whereas complex rule-based targets tracked across time evoked a P3b-like global ERP response which generalised across a late time window. Converging results from temporal cross-modality decoding analyses across different datasets, we reveal that global ERP responses are sustained in a supramodal higher-order network whereas local ERP responses canonically thought to rely on modality-specific regions evolve into short-lived supramodal activity. Taken together, our findings demonstrate that cortical organisation largely follows a gradient in which short-lived modality-specific as well as supramodal processes dominate local responses whereas higher-order processes encode temporally extended abstract supramodal information fed forward from modality-specific cortices. Sensory target detection at local and global timescales reveals a hierarchy of supramodal dynamics in the human cortex

**Significance statement:** Each sense supports a cortical hierarchy of processes tracking deviant sensory events at multiple timescales. Conflicting evidence produced a lively debate around which of these processes are supramodal. Here, we manipulated the temporal complexity of auditory, tactile, and visual targets to determine whether cortical local and global ERP responses to sensory targets share cortical dynamics between the senses. Using temporal cross-decoding, we found that temporally complex targets elicit a supramodal sustained response. Conversely, local responses to temporally confined targets typically considered modality-specific rely on early short-lived supramodal activation. Our finding provides evidence for a supramodal gradient supporting sensory target detection in the cortex, with implications for multiple fields in which these responses are studied (such as predictive coding, consciousness, and attention).

## Introduction

The ability to detect deviant sensory events in a stream of predictable stimuli is crucial for adaptive behaviour. To enable this, each sense relies on a dedicated temporospatial hierarchy of cortices spanning from primary sensory to associative and frontal areas in which successive levels encode increasingly abstract sensory information (Çatal, Gomez-Pilar, & Northoff, 2022; de Lange, Heilbron, & Kok, 2018; Golesorkhi, Gomez-Pilar, Tumati, Fraser, & Northoff, 2021; Ito, Hearne, & Cole, 2020; Kiebel, Daunizeau, & Friston, 2008; Raut, Snyder, & Raichle, 2020; Taylor, Hobbs, Burroni, & Siegelmann, 2015; Wengler, Goldberg, Chahine, & Horga, 2020).

Sensory targets are followed by two cortical responses which can be located on successive levels of the cortical hierarchy based on their temporal and cognitive properties: the Mismatch Negativity (MMN) and the P300 (P3a/P3b) complex. The MMN is an early local negativity associated with temporally proximal sensory change detection in different sensory modalities

(Allen et al., 2016; Czigler et al., 2006; Mäntysalo & Näätänen, 1987). Canonically considered pre-attentive (Tiitinen, May, Reinikainen, & Näätänen, 1994), the MMN is modulated by attention but resists distraction (Auksztulewicz & Friston, 2015; Chennu et al., 2013). The P3b is a late distributed temporally extended positivity indexing complex targets across sensory modalities (Pegado et al., 2010; Polich, 2007; Yamaguchi & Knight, 1991b, 1991a). Unlike the MMN, the P3b requires attention and memory to track the sensory context surrounding targets (Katayama & Polich, 1998; Polich, 2007; Squires, Petuchowski, Wickens, & Donchin, 1977). Based on these properties, the MMN and P3b can be placed in lower-order and higher-order levels in the cortical hierarchy respectively (Bekinschtein et al., 2009; Chao, Takaura, Wang, Fujii, & Dehaene, 2018; Wacongne et al., 2011).

Earlier work suggests that the prefrontal cortex processes complex, temporally extended targets while the detection of simple, temporally confined targets largely recruits modality-specific areas (Bekinschtein et al., 2009; Cornella, Leung, Grimm, & Escera, 2012; Donner et al., 2000; Ester, Serences, & Awh, 2009; Golesorkhi et al., 2021; Maekawa et al., 2005; Miller, 2009; Wacongne et al., 2011; Wolff et al., 2022). Nevertheless, considerable debate revolves around the extent to which local and global ERP responses to sensory targets rely on supramodal activation patterns. Various studies found the P3b to originate in supramodal, but also modality-specific frontoparietal sources (Dreo, Attia, Pirtošek, & Repovš, 2017; Halgren et al., 1995; Halgren, Marinkovic, & Chauvel, 1998; Katayama & Polich, 1998; Walz et al., 2013). Primary sensory and inferior frontal sources generate the MMN in different sensory modalities (Akatsuka, Wasaka, Nakata, Kida, & Kakigi, 2007; Garrido, Kilner, Stephan, & Friston, 2009a, 2009b; Näätänen, Simpson, & Loveless, 1982; Ostwald et al., 2012; Pazo-Alvarez, Cadaveira, & Amenedo, 2003; Shen, Smyk, Meltzoff, & Marshall, 2018). Evidence investigating supramodal contributions to the MMN is inconclusive (Chang, Seth, & Roseboom, 2017; Mariola, Baykova, Chang, Seth, & Roseboom, 2019), leaving the question open whether local sensory mismatch responses share common neural signatures between the senses.

We hypothesised that global top-down-driven ERP responses to complex targets might share neural signatures between the senses. In contrast, local bottom-up-driven ERP responses to simple targets might be supported by early localised modality-specific activity with only few supramodal contributions. Our approach exploits differences in the susceptibility of electrophysiological responses to bottom-up and top-down variables to dissociate their neural dynamics in different levels of the cortical hierarchy. Based on earlier work elucidating local and global cortical signalling in the auditory hierarchy (Bekinschtein et al., 2009; Chennu et al., 2016; Phillips et al., 2016; Sitt et al., 2014; Wacongne et al., 2011). we use multisensory versions of a hierarchical oddball paradigm (“local-global paradigm”) in which local and global irregularities in the sensory environment elicit P3b-like global responses as well as MMN-like local responses (Shirazibeheshti et al., 2018). The local-global paradigm achieves this by manipulating the complexity of the sensory context in which a stimulus change appears, thereby mapping cortical and perceptual local-global hierarchies onto each other (Northoff, Wainio-Theberge, & Evers, 2020).

## Materials and methods

### Participants

We developed two multisensory versions of the local-global paradigm. In the bimodal version, separate blocks of somatosensory or auditory expectation violations were presented. In addition to purely visual, somatosensory and auditory blocks, the trimodal version of the paradigm encompassed blocks in which trials combined inputs from two different sensory modalities which could either be visual, auditory or somatosensory. Only individuals with no history of neurological or psychiatric conditions and no tactile and auditory impairment were recruited into both studies. In addition, individuals with visual impairments were excluded from participation in the trimodal study. All participants gave written and informed consent. Data for the bimodal local-global paradigm were collected at the University of Cambridge and obtained ethical approval from the Department of Psychology (CPREC 2014.25). For the bimodal study, we invited individuals aged 18-35 to participate through the SONA participant database at the Department of Psychology. We recruited 30 individuals (15/15 female/male, mean+-STD age is 24.57(+-4.52)) who were paid £10 per hour for a duration of 3-3.5 hours. The trimodal task was performed at the Centre Nationale de la Recherche Scientifique (CNRS) in France. Participants for this study were invited through the CNRS RISC system. Only individuals aged 18-80 were asked to participate in the trimodal study. Of 54 participants (35 female, mean(±STD) age: 25.20(±4.10) years) who participated in the trimodal study, 7 were excluded due to a recording error. Participants in the trimodal study were paid €40 for their effort.

### Materials

In the trimodal study, auditory stimulation was applied using Etymotic noise-isolating insert earphones. Eccentric Rotating Mass motors controlled by two Texas Instruments DRV2605 haptic drivers were employed to deliver vibrotactile stimulation to the wrist. Two independent 8×8 LED matrices implemented in virtual reality goggles were used to apply visual inputs isolating visual hemifields. The setup was controlled using an Arduino Zero board. The bimodal study used auditory inputs generated by mixing three sinusoidal signals of either 500, 1000, and 2000 Hz (tone type A) or 350, 700, and 1400 Hz (tone type B) in Matlab R2016 based on (Chennu et al., 2013) and applied using EARTONE 3A insert earphones. Tactile stimulation was delivered using a custom-made device which applies mechanical pins to the fingertip with a Saia-Burgess 12.7 mm stroke, 12 v, 4 W DC push-action solenoid with 0.3-0.6 N force, and no nominal delay after current onset) controlled by an Arduino Mega board.

### Experimental design

We designed two multisensory variants of the local-global oddball paradigm depicted in Figure 1. In this paradigm, expectations about sensory inputs are violated either locally within trials or globally between trials. Here, trials were composed of five stimuli with a stimulus duration of 50 ms and a stimulus onset interval of 150 ms. Each trial consisted of four identical ipsilateral stimuli followed by a deviant contralateral stimulus (locally deviant trials) or another ipsilateral stimulus (locally standard trials). Contrasting locally deviant and standard trials reveals a MMN-like amplitude difference referred to as local ERP response in a time window between 50-250 ms after onset of the last stimulus in a trial. Global violations of sensory expectations are achieved when a frequently presented trial type is occasionally interspersed with a different trial type. A comparison of frequent (globally standard) trials and rare (globally deviant) trials uncovers a global ERP response which manifests as a late distributed P3b-like positive wave (Bekinschtein et al., 2009; Chennu et al., 2013, 2016; Phillips et al., 2016; Sitt et al., 2014; Wacongne et al., 2011).

**Figure 1.**
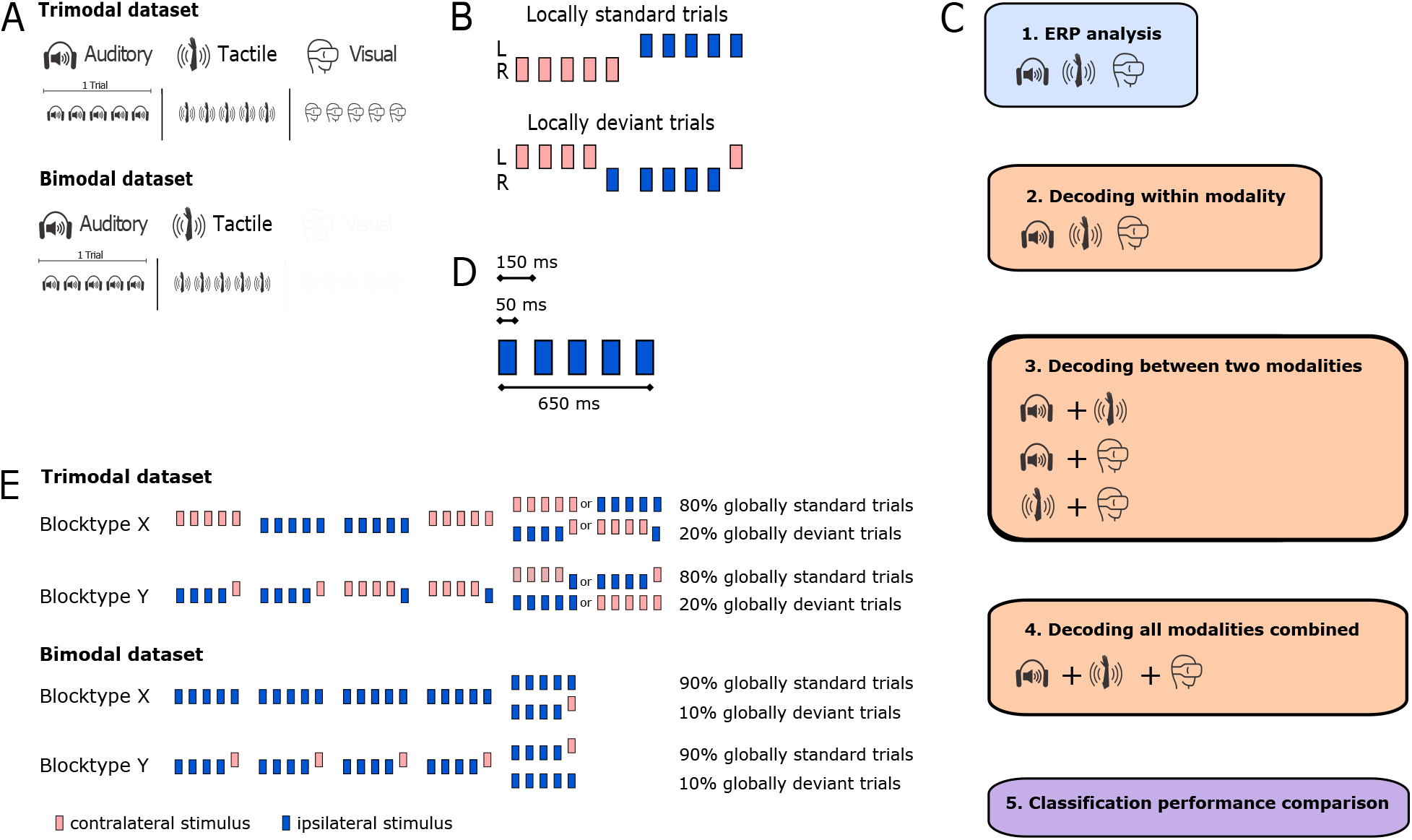
Experimental design. Each trial consists of 5 stimuli sampled from one sensory modality. Stimuli could either be auditory, visual or tactile in the trimodal study or auditory and tactile in the bimodal study. For each block, trials sampled from specific sensory modality are presented (A). *Trial design setting up the local contrast between deviant and standard stimuli at trial end*: Standard trials consisted of five identical stimuli applied to the same body hemisphere. Deviant trials were composed of four identical ipsilateral stimuli followed by a contralateral stimulus. In the trimodal study, sensory stimulation started on the left side in 50% of trials for each block. In the bimodal study, trials in 50% of blocks started on the left (B). We first analysed ERP time courses to obtain local and global ERP responses for each contrast. Then, we performed a series of three main temporal decoding analyses for each dataset: First, we decoded the temporal evolution of each local and global ERP response within a sensory modality. We trained and classifiers on one modality to test them on another for each modality pair. In a further step, we trained and tested classifiers on a combination of trial from all sensory modalities for local and global ERP responses separately. Finally, we performed comparisons of classification performance between local and global ERP responses (C). Each stimulus was presented for 50 ms with an inter-stimulus interval of 150 ms. Trials were presented with a jittered inter-trial interval of 1450-1650 ms in the bimodal study and a fixed inter-trial interval of 1400 ms for the trimodal study (D). *Block design setting up the global contrast using nested stimulus groups*: In blocktype X, locally standard trials dominate the input stream and locally deviant trials occur only rarely at a global level. In blocktype Y, this pattern is inverted (E).

During each study, two block types were presented. The bimodal study consisted of 8 auditory and somatosensory blocks. Standard stimuli in 50% of blocks were presented on the left side and on the right side in the remaining blocks. Each block consisted of 78% globally standard trials and 22% globally deviant trials. In block type X, locally standard trials were occasionally interrupted by 22 % locally deviant trials which could equally likely be a locally deviant trial in which only the laterality or both laterality and stimulus type were varied. In block type Y, a stream of locally deviant trials in which the last stimulus was applied to the contralateral body hemisphere was occasionally interrupted by locally standard trials or locally deviant trials in which the last stimulus differed in laterality and stimulus type. In the trimodal study, blocks were composed of 80% globally standard and 20% globally deviant trials. Trials randomly started on the left or right side within each block with 50% probability. In blocktype X, locally standard trials consisting of five identical stimuli were interspersed with locally deviant trials in which the last stimulus was applied to the contralateral hemisphere. Blocktype Y consisted of a sequence of locally deviant trials occasionally interrupted by locally standard trials. Blocks in both tasks started with a habituation phase in which the globally standard trial was repeated to establish an expectation of globally recurring stimulus patterns. We presented 24 repetitions of the globally standard in the trimodal study and 15 repetitions in the bimodal study.

In the bimodal study, somatosensory locally standard trials consisted of five touches ipsilaterally applied to the index finger. We introduced two types of locally deviant trials in which the last stimulus in a trial was applied to the contralateral index finger or contralateral middle finger. In auditory blocks, locally standard trials presented as five identical sounds. Local deviations were introduced by varying either only the laterality of the ear which received the last sound in a trial or both the laterality and pitch of the last sound in a trial. The latter locally deviant trial type was globally deviant type in each block type and the former locally deviant trial type was globally standard in block type Y and deviant in block type X. The bimodal study consisted of 8 auditory and 8 somatosensory blocks. The block order was pseudo-randomised so that the experiment started with somatosensory block type Y followed by somatosensory block type X, no more than two consecutive blocks were presented in the same sensory modality, and each half of the experiment contained equal proportions of somatosensory and auditory blocks. Each block consisted of 158-160 trials and lasted ∼4.5 min. Inter-trial intervals were randomly sampled from a uniform distribution between 800-1000 ms in steps of 50 ms. In each block, 30-34 globally deviant trials were embedded in a sequence of 112 globally standard trials and both deviant types occurred in equal proportions. Each globally deviant trial was preceded either by 2, 3, 4 or 5 globally standard trials in equal proportions. Participants were exposed to white noise during somatosensory blocks to avoid auditory cues from the tactile stimulation device.

The trimodal study contained blocks with exclusively auditory, visual or somatosensory stimulation. Participants in this study also underwent multisensory blocks in which expectations violations require to converge inputs from two of these sensory modalities and which were not further analysed. Local deviations were introduced by varying the laterality of the last stimulus in a trial. 50% of trials applied standard stimuli to the left hemisphere (ear, visual hemifield, or wrist) and the last stimulus to the right hemisphere (and vice versa). Participants underwent three experimental sessions with a total duration of 20 min (4.5 min per session). The remaining two sessions applied crossmodal stimulation and analyses were not included in this paper. 16 participants were presented with somatosensory blocks, 15 participants with auditory blocks and 16 participants with visual blocks. Each block type was presented twice per session in a fixed order: X-Y-X-Y. 31 (∼20%) trials included in each block were globally deviant. Each globally deviant trial was preceded by 3, 4, or 5 globally standard trials. To ensure that participants attend to the global regularity in sensory stimulation patterns, we instructed them to count the number of deviant stimulus groups occurring in a stimulus stream and report the number after each block. Blocks in which participants deviated from the true count by more than two were repeated.

### Statistical analysis

EEG data acquisition was performed using a Net Amps 300 amplifier with an Electrical Geodesics 256-channel high density EEG net at the ICM in Paris for the trimodal study and an Electrical Geodesics 128-channel high density EEG net at the Department of Psychology, University of Cambridge for the bimodal study. We performed EEG data preprocessing using EEGLAB 2019 in Matlab 2019b. In a first step, data were down-sampled to 250 Hz, filtered between 0.5-30 Hz and epoched with respect to the onset of the last stimulus in a trial. Habituation trials at the start of each block were removed. Having removed electrodes placed on the neck and cheek which record mostly muscle artefacts, we retained 91 electrodes in the bimodal data set and 175 electrodes in the trimodal data set for further analysis. We performed baseline removal using a window of 100 ms before epoch onset. Noisy trials (with a variance of >350) and channels (with a variance of >500) were temporarily removed using a semi-automated procedure. We removed artefacts resulting from sweat, eye and muscle movements using independent component analysis. Ultimately, we removed the remaining artefacts using trial-wise interpolation.

We performed cluster-based permutation analyses to test for differences in ERP amplitude time courses. We used the Common Average as a reference and performed a baseline correction in a time window of 100 ms before onset of the last stimulus in a trial. For each condition pair with unequal trial numbers, trials in the condition with a higher trial number were randomly deselected until the number of trials in both conditions was equal. Cluster-based permutation uses Monte Carlo partitioning to obtain a cluster-level t-statistic. To perform Monte Carlo partitioning, data are pooled and randomly divided into two new data sets of equal size. We performed two-sided t-tests on the subject averages time-channel pairs and retained only t-values with p < 0.05. Spatiotemporally adjacent t-values were summarised and the largest cluster-level summarised t-value was identified. Having performed this procedure 3000 times, we determined the p-value corresponding to the proportion of maximal cluster-level t-values larger than the observed t-value in the original comparison. Conditions were deemed to be different if p < 0.05 (Oostenveld, Fries, Maris, & Schoffelen, 2011).

We applied temporal decoding to examine whether two contrasts rely on similar cortical signatures. Temporal decoding is a machine learning procedure which assesses whether a classifier trained to discriminate two trial types at one time point will generalise to the remaining time points in a sample. We applied a bootstrapping procedure in which 5 trials were randomly sampled from each dataset until we reached 540 epoch averages for each trial type (deviant/standard) respectively. This procedure was repeated 50 times. Classification performance was assessed on each of the resulting datasets. To perform temporal decoding within a sensory modality, we used a 5-fold stratified cross-validation procedure in which linear support vector machines were trained to optimally separate standard and deviant trials on 4/5 of the data set and tested on the remaining data. For temporal decoding between sensory modalities, we fitted classifiers using training and testing datasets from two different sensory modalities. For each condition pair, we trained and tested classifiers on every time point in a time window of 600 ms after onset of the last stimulus in a trial. Classification performance was assessed using the Area Under the Curve Receiver Operating Characteristic (AUC-ROC) which is a non-parametric criterion-free measure of separability. This procedure results in a training time vs testing time temporal generalisation matrix with AUC-ROC classification scores as cells. To identify adjacent AUC-ROC scores which differ from chance, we performed a Monte Carlo cluster-based permutation analysis with 1024 random partitions on classification score averages from each bootstrapped dataset and applied two-tailed paired t-tests to identify clusters of AUC-ROC values (p < 0.05) which differ from chance (King & Dehaene, 2014; King, Gramfort, Schurger, Naccache, & Dehaene, 2014).

## Results

### Hierarchically nested sensory deviations elicit local and global ERP responses in different sensory modalities

We investigated commonalities in cortical responses to rule-based global and feature-based local sensory deviations using multisensory versions of the local-global paradigm.

In a first step, we established that hierarchical manipulations of sensory context elicit a MMN-like local ERP response and a P3b-like global ERP response in the auditory, somatosensory and visual modality in both experiments. To that end, we performed cluster-based permutation of ERP amplitude time courses (Maris & Oostenveld, 2007) displayed in Figure 2. Replicating earlier findings (Bekinschtein et al., 2009; Chennu et al., 2013), the auditory local ERP response in the bimodal paradigm manifested as a frontotemporal bipolar two-peak difference wave (cluster t = −12000, p < 0.001) and a one-peak difference wave in the trimodal paradigm (cluster t = −5396.6, p < 0.001). A somatosensory local ERP response emerged as a central negativity between ∼50-150 ms in the bimodal paradigm (cluster t = −752.31, p < 0.001) and as a two-peak negativity between ∼100-350 ms the trimodal paradigm (cluster t = −3298.4, p < 0.001). We also identified a visual local ERP response as a negativity in a mid-range time window between ∼100-350 ms (cluster t = −6178.9, p < 0.001) shown in Figure 2.

**Figure 2.**
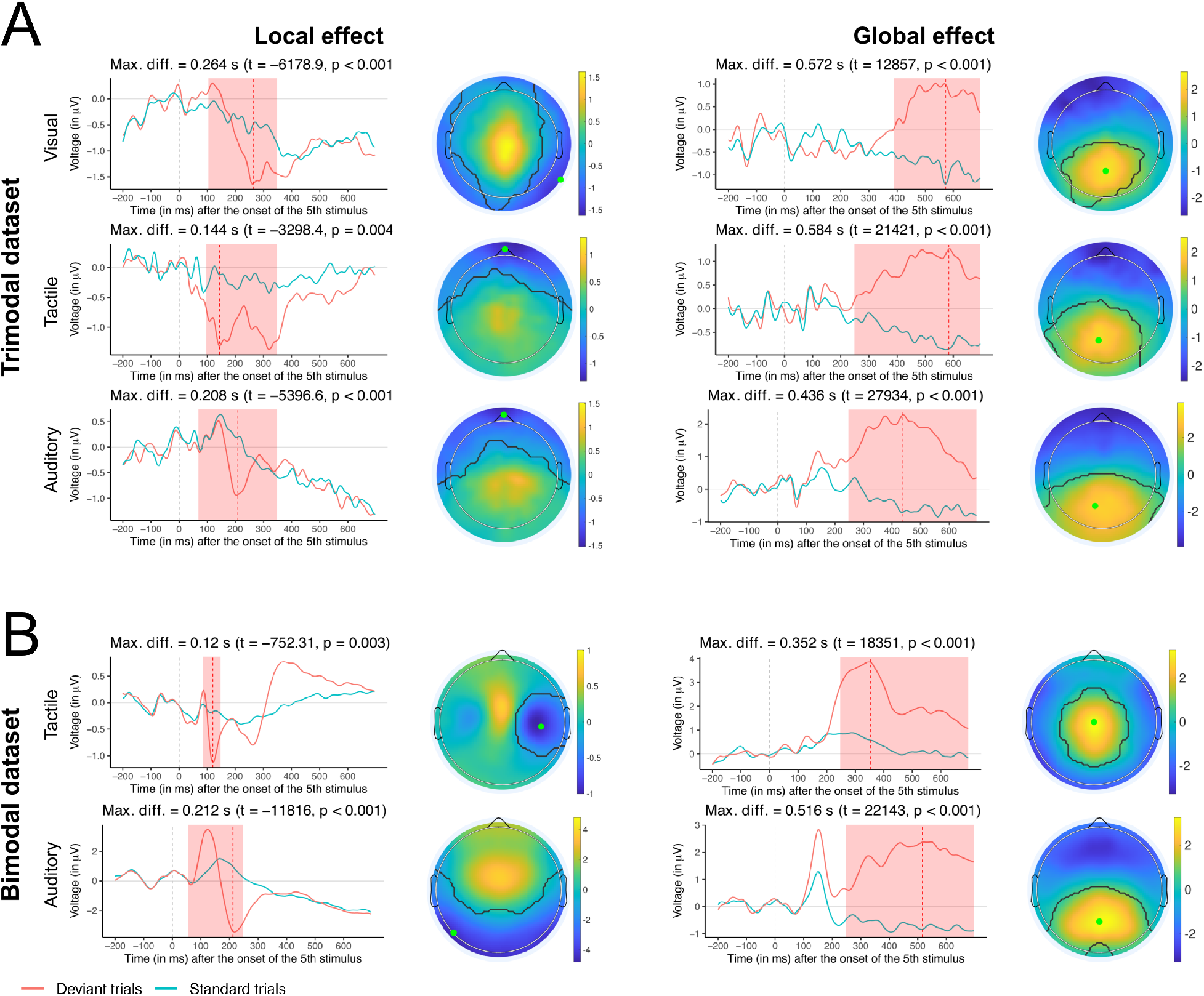
Experimental design and ERP results. Cluster-based permutation test results for ERP amplitude differences showing that a local and global ERP response can be obtained in different sensory modalities for the trimodal (A) and the bimodal study (B). On the left, each panel shows ERP amplitude time courses of the corresponding deviant-standard condition pair. Next to the ERP voltage time course plot, the corresponding topographical map is shown. Time periods in which a cluster-based permutation test identified a difference between both conditions are highlighted in red in the ERP time course plot and delineated with a black line on the topographical map. The time point at which the difference between both conditions is maximal is delineated with a dotted red line in the ERP amplitude time course plot. The electrode at which this difference was obtained is marked with an orange dot on the topographical map.

A comparison of globally deviant and standard trials revealed a positive difference wave in a time window from ∼250 ms until the end of the trial regardless of sensory modality (with some shifts in onset of the effect). In the bimodal (cluster t = 22143, p < 0.001) and trimodal study (cluster t = 27934, p < 0.001), we revealed an auditory global ERP response as a positive deflection with a posterior distribution. A somatosensory global ERP response presented with a similar posterior distribution in both the bimodal (cluster t = 18351, p < 0.001) and trimodal study (cluster t = 21421, p < 0.001). Ultimately, a visual global ERP response emerged as a positive difference wave in a relatively late time window from ∼400 ms until the end of the trial (cluster t = 12857, p < 0.001) shown in Figure 2. These results show that complex targets which require the conscious tracking of sensory patterns across time elicit a global ERP response in different sensory modalities. Our finding that the global ERP response manifests as a large, late and posterior positive deflection regardless of sensory modality complements previous studies which characterise the related P300 as a late positivity (Bekinschtein et al., 2009; Bledowski et al., 2004; Chennu et al., 2013; Walz et al., 2013), Taken together, our results show that the functional dissociation of local and global ERP responses is a supramodal property of target detection systems in different sensory domains.

### Cortical responses to rule-based but not feature-based sensory targets are sustained across time in each sensory modality

Previous work has shown that functional differences between cortical responses to auditory rule-based and feature-based targets are reflected in the extent to which they are maintained in auditory networks. Cortical activation patterns in response to auditory rule-based targets are sustained in time, whereas feature-based auditory deviations decay quickly (King & Dehaene, 2014). Is the temporal evolution of cortical target detection responses a property common to different sensory domains? We examined whether global ERP responses are linked to a sustained cortical activation pattern whereas local ERP responses are supported by short-lived activity regardless of sensory modality. We employed temporal decoding to characterise neural activation patterns elicited by local and global ERP responses in different sensory modalities. In short, temporal decoding is a machine learning approach used to characterise the temporal evolution of cortical activation patterns linked to sensory events. A classifier trained at a time point t is not only tested at t but at all other remaining time points. This leads to a temporal generalisation matrix of classification performance scores. The shape of the temporal generalisation matrix offers insights into the temporal dynamic of cognitive operations and their cortical generators (Dehaene & King, 2016).

We observed that local ERP responses could be decoded along the diagonal in a mid-range time window for each sensory modality (Figure 3). Across different contrasts, local ERP responses were found to decay quickly. This finding is mostly consistent with a serial activation of different cortices dedicated to the sensory modality in which the local ERP response was applied (King et al., 2014). The visual local ERP response was maintained briefly between 200-400 ms (mean AUC = 0.51 ± 0.03, max. AUC = 0.61 at 268 ms training time and 264 ms testing time, mean cluster t = 3.8, p < 0.05). In the trimodal study, the somatosensory local ERP response (mean AUC = 0.51 ± 0.02, max. AUC = 0.58 at 212 ms training time and 216 ms testing time, mean cluster t = 2.96, p < 0.05) and the auditory local ERP response (mean AUC = 0.52 ± 0.03, max. AUC = 0.67 at 220 ms training time and 212 ms testing time, mean cluster t = 4.52, p < 0.05) were maintained for ∼100 ms from ∼200 ms. In the bimodal study, the somatosensory effect was best decoded between 200-300 ms (mean AUC = 0.52 ± 0.03, max. AUC = 0.63 at 208 ms training time and 208 ms testing time, mean cluster t = 4.47 ± 7.53, p < 0.05). In this study, temporal decoding revealed a classification score matrix consistent with two distinct processes underpinning the auditory local ERP response. From ∼100 ms, the auditory local ERP response can be decoded along the diagonal which suggests a serial propagation of cortical activity along the auditory cortical hierarchy. However, cortical activity is sustained for ∼150 ms from ∼200 ms (mean AUC = 0.51 ± 0.04, max. AUC = 0.71 at 188 ms training time and 188 ms testing time, mean cluster t = 2.11 ± 13.15, p < 0.05). Although classification scores for the auditory and somatosensory local ERP response differed from chance across an extended time window, classifiers performed only slightly better than chance from ∼350 ms for both effects.

**Figure 3.**
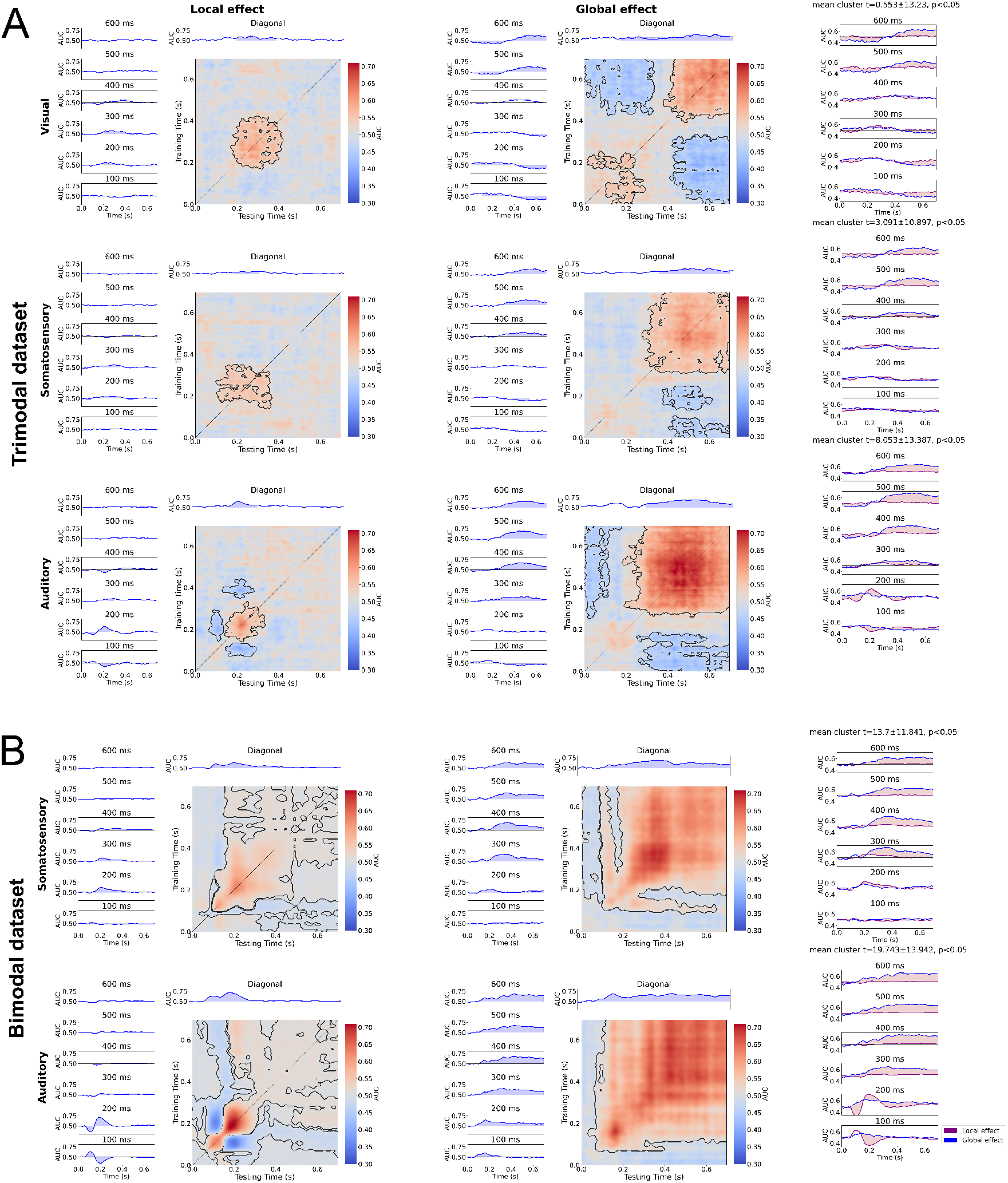
Temporal generalisation analysis within sensory modalities. Panels display temporal generalisation results for the local (left) and global ERP response (right) in the trimodal (A) and bimodal dataset (B). Each panel shows results of a temporal generalisation analysis in which a classifier is trained to distinguish deviant and standard trials at each time point and then tested on all remaining time points in a trial. Classification scores are displayed on a red-to-blue gradient. On top of each matrix, adjacent classification scores different from chance (0.5) in a cluster-based permutation test are highlighted with a black line. Next to each matrix, we show a series of time courses of classification scores produced by classifiers trained at the specified time point. On top of each matrix, we plotted the classification score time course for a decoding procedure in which a classifier is trained and tested on deviant-standard pairs at the same time point (corresponding to the matrix diagonal). Classification scores different from chance are highlighted with a blue shade. On the right side, we show results of a cluster-based permutation test contrasting classification score matrices of the corresponding local and global ERP response. Each subplot shows the classification score time courses of a classifier trained to separate global deviant-standard pairs at the specified time point and tested across the remaining time window, and its local counterpart. Time periods in which local and global ERP response classification time courses differ are highlighted in red shade.

We found that classifiers trained to distinguish globally deviant and standard trials from ∼200 ms generalised across other time samples in the remaining trial window regardless of which sensory modality was tested (Figure 3). This decoding procedure led to a rectangular classification score matrix for each comparison. In line with earlier ERP time course comparison results (Figure 2), the visual global ERP response manifested relatively late from ∼400 ms (mean AUC = 0.51 ± 0.06, max. AUC = 0.66 at 516 ms training time and 520 ms testing time, mean cluster t = 3.19, p < 0.05). In the trimodal study, the somatosensory global ERP response appeared from ∼350 ms ((mean AUC = 0.52 ± 0.05, max. AUC = 0.64 at 512 ms training time and 508 ms testing time, mean cluster t = 5.56, p < 0.05)) and the auditory global ERP response from ∼200 ms (mean AUC = 0.54 ± 0.07, max. AUC = 0.69 at 532 ms training time and 524 ms testing time, mean cluster t = 12.5, p < 0.05). In the bimodal study, temporal generalisation was found relatively early from ∼150 ms for both the somatosensory (mean AUC = 0.56 ± 0.06, max. AUC = 0.7 at 372 ms training time and 344 ms testing time, mean cluster t = 19.32 ± 19.07, p < 0.05) and auditory global ERP response (mean AUC = 0.6 ± 0.05, max. AUC = 0.7 at 432 ms training time and 428 ms testing time, mean cluster t = 33.06 ± 17.56, p < 0.05). Cluster-based permutation tests comparing differences in classification scores between the local and global ERP response show that the global ERP response generalises in a late time window whereas the local ERP response does not (Figure 3).

Collectively, our results indicate that cortical activation associated with the global ERP response starts no earlier than ∼150 ms after onset of the last stimulus in a trial and is sustained over time until the trial ends. However, we observed some shifts in the onset latency between different global ERP responses with the visual global ERP response not appearing before 400 ms. Despite these shifts, this pattern suggests that a single cortical system is active in that time window (King & Dehaene, 2014). This finding leaves the question open whether this system is dedicated to a specific sensory modality or shared between different senses.

### Supramodal activation is sustained for global but decays for local ERP responses

Here we provide evidence for the hypothesis that cortical hierarchies dedicated to each sense are organised along a gradient of supramodality. Building up on our finding that global ERP responses are associated with a sustained late cortical activation pattern, we further demonstrate that this sustained pattern is shared between different sensory modalities. We also show that local ERP responses in different sensory modalities rely on few, if any, common cortical signatures in comparison to global ERP responses.

Temporal decoding was employed to examine whether cortical responses share neural dynamics between sensory modalities. To identify common activity patterns linked to evoked responses, temporal generalisation analysis can be applied in two ways: Classifiers can be trained to separate a deviant-standard condition pair in a target sensory modality at a time point t and tested on deviant-standard condition pairs in a different sensory modality across all time points in a trial. Alternatively, a classifier can be trained to separate deviant-standard condition pairs when they are pooled across all sensory modalities separately for global and local ERP responses. We used a 5-fold stratified cross-validation approach in which classifiers were trained on 4/5 of the data and tested on the remaining 1/5 for both analyses (see Methods for details).

We initially trained a classifier to separate globally deviant from standard trials when trials for each condition pair are pooled across sensory modalities. This procedure revealed temporal generalisation in a late time window across different global ERP responses in both datasets (Figure 4). We provide evidence for shared activity supporting the auditory and visual global ERP response from ∼400 ms regardless of whether classifiers were trained on the visual and tested on the auditory contrast (mean AUC = 0.52 ± 0.07, max. AUC = 0.66 at 452 ms training time and 572 ms testing time, mean cluster t = 4.62, p < 0.05) or vice versa (mean AUC = 0.51 ± 0.06, max. AUC = 0.65 at 676 ms training time and 496 ms testing time, mean cluster t = 3.37, p < 0.05). A similar pattern was found for somatosensory and visual effects when classifiers were trained on visual and tested on somatosensory deviant-standard pairs (mean AUC = 0.51 ± 0.05, max. AUC = 0.64 at 540 ms training time and 464 ms testing time, mean cluster t = 3.2, p < 0.05) or vice versa (mean AUC = 0.51 ± 0.07, max. AUC = 0.68 at 580 ms training time and 540 ms testing time, mean cluster t = 2.94, p < 0.05). Shared activity between the auditory and somatosensory global ERP response was found from ∼350 ms when classifiers were trained on the auditory and tested on the somatosensory contrast (mean AUC = 0.53 ± 0.05, max. AUC = 0.64 at 424 ms training time and 580 ms testing time, mean cluster t = 10.21, p < 0.05) or vice versa (mean AUC = 0.53 ± 0.06, max. AUC = 0.69 at 576 ms training time and 432 ms testing time, mean cluster t = 10.26, p < 0.05). This finding was replicated in the bimodal dataset in a more extensive time window from ∼150 ms when classifiers were trained on the somatosensory and tested on the auditory effect (mean AUC = 0.57 ± 0.06, max. AUC = 0.67 at 188 ms training time and 160 ms testing time, mean cluster t = 22.26 ± 19.85, p < 0.05) and vice versa (mean AUC = 0.57 ± 0.05, max. AUC = 0.66 at 292 ms training time and 348 ms testing time, mean cluster t = 24.75 ± 17.7, p < 0.05). Taken together, these findings show that there is shared activation between global ERP responses regardless of which sensory modality is used for training and testing. The global ERP response consistently manifests in a rectangular classification score matrix which suggests that common activation is maintained in cortical networks (with some temporal shifts in onset times).

**Figure 4.**
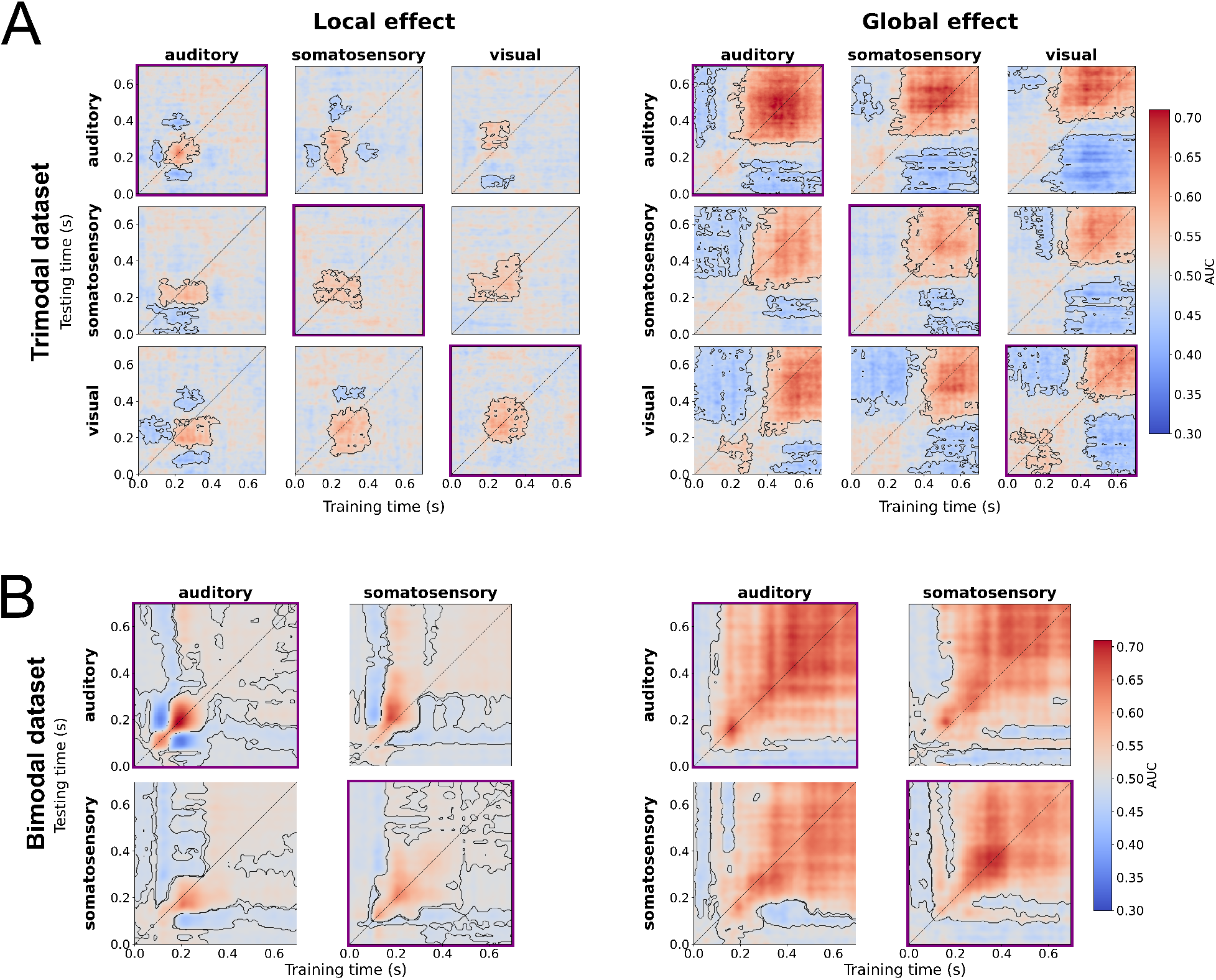
Temporal generalisation analysis within and between sensory modalities. Each panel shows results from a series of temporal generalisation analyses decoding the local ERP response (left) and the global ERP response (right) in the trimodal dataset (A) and bimodal dataset (B). Each matrix shows results of a temporal generalisation analysis in which a classifier is trained to distinguish deviant and standard trials at each time point in the training modality and then tested on all remaining time points in a trial in the testing modality. For each experiment, this leads to an n-by-n matrix with n being the number of sensory modalities tested in a study (3 for the trimodal and 2 for the bimodal study). Temporal generalisation matrices placed along the diagonal of a panel show results from a decoding analysis performed within a sensory modality and are highlighted with a purple frame. The remaining temporal generalisation matrices show results from a decoding procedure in which a classifier is trained to distinguish deviant and standard trials corresponding to the sensory modality indicated next to each row and tested on the sensory modality corresponding to the column label. AUC-ROC classification scores are shown on a red-to-blue gradient. Classification score clusters which were found to be different from chance in a cluster-based permutation test are delineated with a black line. 0 ms describes the onset of the last stimulus in a trial.

Interestingly, temporal decoding of local ERP responses revealed short-lived temporal generalisation around ∼200 ms in both datasets. In the trimodal dataset, short-lived shared representations from ∼200-350 ms were found to be associated with local ERP responses across all comparisons: auditory to somatosensory (mean AUC = 0.51 ± 0.03, max. AUC = 0.58 at 204 ms training time and 252 ms testing time, mean cluster t = 2.04, p < 0.05), somatosensory to auditory (mean AUC = 0.5 ± 0.03, max. AUC = 0.62 at 216 ms training time and 200 ms testing time, mean cluster t = 1.01, p < 0.05), auditory to visual (mean AUC = 0.5 ± 0.03, max. AUC = 0.62 at 176 ms training time and 268 ms testing time, mean cluster t = −0.57, p < 0.05), visual to auditory (mean AUC = 0.52 ± 0.02, max. AUC = 0.59 at 264 ms training time and 192 ms testing time, mean cluster t = 4.64, p < 0.05), somatosensory to visual (mean AUC = 0.5 ± 0.03, max. AUC = 0.6 at 208 ms training time and 260 ms testing time, mean cluster t = 1.34, p < 0.05) and visual to somatosensory (mean AUC = 0.5 ± 0.02, max. AUC = 0.55 at 260 ms training time and 216 ms testing time, mean cluster t = −0.93, p < 0.05)). In the bimodal dataset, we found evidence for shared representations from ∼200 ms when classifiers were trained on somatosensory and tested on auditory deviant-standard pairs (mean AUC = 0.52 ± 0.03, max. AUC = 0.68 at 208 ms training time and 176 ms testing time, mean cluster t = 4.9 ± 9.02, p < 0.05) and vice versa (mean AUC = 0.51 ± 0.02, max. AUC = 0.64 at 172 ms training time and 208 ms testing time, mean cluster t = 2.99 ± 8.74, p < 0.05).

Our result demonstrates that local ERP responses are supported by supramodal transient activation patterns starting from ∼200 ms. Although our finding is compatible with the idea that local ERP responses are processed in cortical hierarchies dedicated to the sensory modality in which the stimulation is applied, we provide evidence for some overlap in higher-order or associative regions between these hierarchies. In sum, these results support the hypothesis that global ERP responses share sustained neural activation patterns between the senses while local ERP responses share fewer, if any, activity (Figure 4).

To corroborate these findings, we use temporal decoding to examine the temporal evolution of cortical activity when global deviant-standard pairs and local deviant-standard pairs are each pooled across sensory modalities. Our results show that the local ERP response is associated with some temporal generalisation from ∼180-250 ms in both the bimodal (mean AUC = 0.51 ± 0.02, max. AUC = 0.65 at 204 ms training time and 204 ms testing time, mean cluster t = 2.75 ± 8.43, p < 0.05)and the trimodal dataset (mean AUC = 0.51 ± 0.03, max. AUC = 0.57 at 220 ms training time and 220 ms testing time, mean cluster t = 3.87, p < 0.05). For local ERP responses, our result is consistent with the involvement of a series of modality-specific neural generators in early levels of the cortical hierarchy and a contribution of supramodal regions in later stages. Again, temporal decoding revealed sustained shared activation starting from ∼150 ms until trial end for the global ERP response in the bimodal dataset (mean AUC = 0.579 ± 0.057, max. AUC = 0.667 at 340 ms training time and 336 ms testing time). In the trimodal dataset, shared activation between global ERP responses extended across the complete time window. We found evidence for sustained activity from ∼350 until trial end. Our results also reveal a rectangular classification score cluster which differs slightly but significantly from chance from the onset of the last stimulus in a trial to ∼250 ms (mean AUC = 0.52 ± 0.06, max. AUC = 0.66 at 520 ms training time and 544 ms testing time, mean cluster t = 7.35, p < 0.05). Based on work by King et al. (King & Dehaene, 2014), these temporal generalisation results suggest that there is a common generator for global ERP responses in different sensory modalities. In contrast, multiple neural generators support the local ERP response. For the local ERP response, our evidence shows that these generators are modality-specific in early stages and supramodal in later stages of cortical processing

Finally, we examined differences in decoding strength and cluster size between the local and global ERP response. For that, we compared maximum classification scores as well as the number of AUC scores with decoding performance above chance in clusters identified by the cluster-based permutation test of decoding performance scores drawn from all 14 temporal decoding analyses using Mann-Whitney U tests. Cluster size was enhanced for the global ERP response relative to the local ERP response (U = 2, p < 0.001), which suggests that activation in supramodal networks supporting the global ERP response is sustained in time, whereas supramodal signatures of the local ERP response decay quickly (Figure 5). Peak decoding performance was also found to be larger for the global than the local ERP response (U = 36, p = 0.001), indicating that decoding results for supramodal activation linked to the global ERP response are relatively more informative.

**Figure 5.**
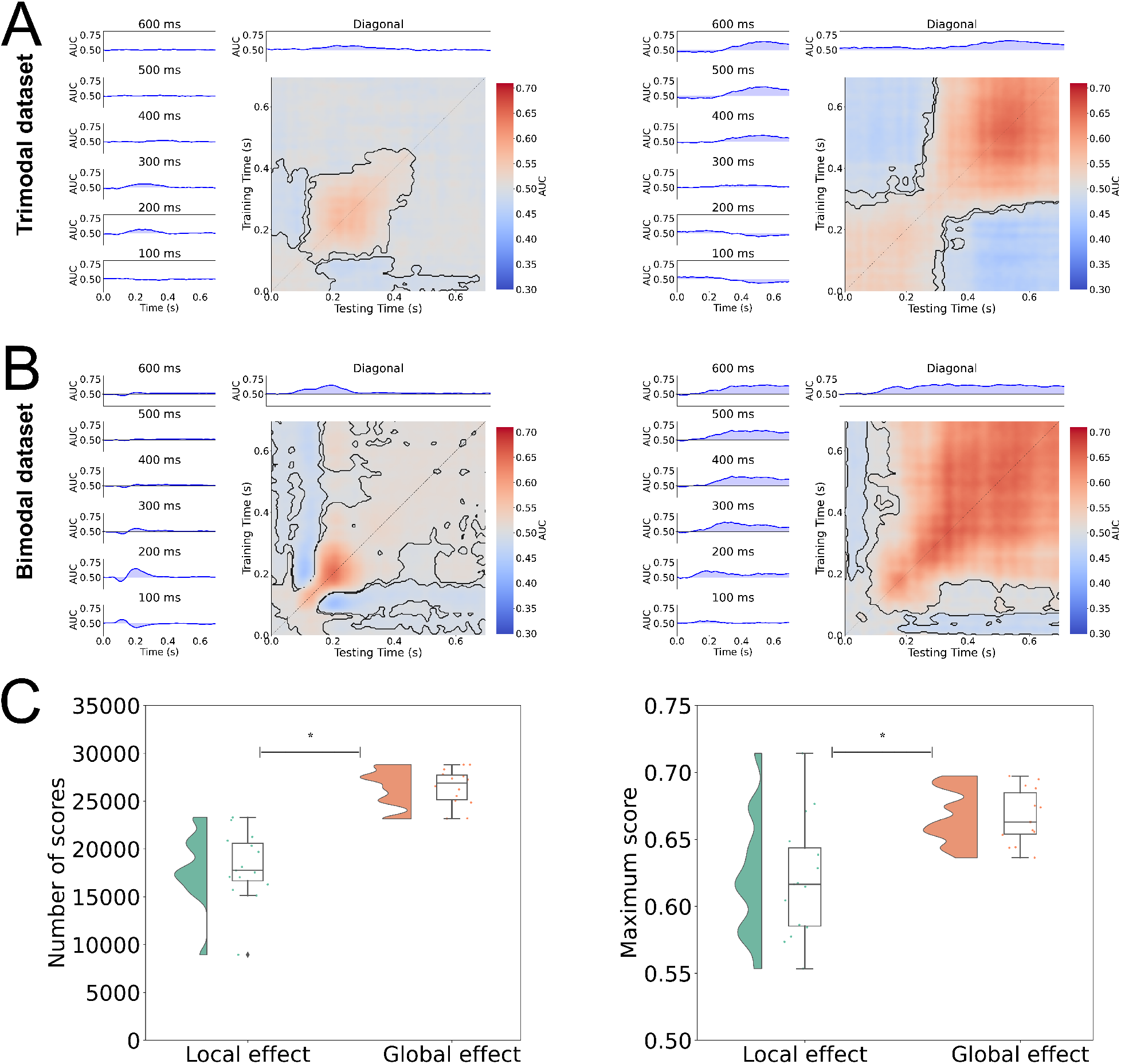
Temporal decoding of combined sensory modalities. In a temporal decoding analysis, we pooled deviant trials and standard trials regardless of sensory modality and tested whether a classifier trained to discriminate deviant and standard trials at a specific time point might generalise to the remaining time points. Each panel shows the resulting matrix of AUC-ROC classification scores for the global (A) and local ERP response (B). In each panel, results for the global ERP response are shown on the right and results for the local ERP response are displayed on the left. In conditions in which cluster-based permutation was performed, clusters which differ from chance are highlighted with a purple horizontal line for classification performance in intervals of 100 ms and green for diagonal classification performance. 0 ms marks the onset of the last stimulus in a trial. Clusters of AUC-ROC scores which differed from chance were delineated with a dotted line. Rain cloud plots supplemented with box-and-whisker plots show the distribution of maximum classification scores (right) and number of classification scores above chance in each cluster (left) drawn from clusters across all 14 temporal decoding analyses for the local and global ERP response. Significant differences between the local and global ERP response were assessed with a Mann-Whitney U Test and highlighted with an asterisk (C).

## Discussion

### Hierarchically nested sensory targets elicit local and global ERP responses across sensory modalities

A long-standing debate in neuroscience revolves around which sensory and perceptual processes are supramodal or modality-specific (Cao, Summerfield, Park, Giordano, & Kayser, 2019; Driver & Noesselt, 2008; Faivre, Filevich, Solovey, Kühn, & Blanke, 2018; Walz et al., 2013). Inspired by the notion that perceptual systems in the cortex are hierarchically organised on a simple-to-complex axis (Dürschmid et al., 2016; Kiebel et al., 2008; Murray, 2004; Rao & Ballard, 1999), we investigated whether cortical responses to sensory targets implemented in successive levels of the cortical hierarchy are ordered along a gradient of supramodality. Across a series of two local-global experiments combining evidence from the somatosensory, visual and auditory modality, we first established that selective hierarchically nested divergences of sensory inputs can trigger an MMN-like local ERP response and a P3b-like global ERP response in different sensory modalities. Most research on the MMN and P3b concentrates on the auditory domain, and comparably less is known about the visual or somatosensory P3b or MMN and their temporal dynamics (Linden et al., 1999; Ostwald et al., 2012). As has been shown for the auditory modality (King et al., 2014), the global ERP response is maintained in higher-order cortical networks while the local ERP response is serially propagated along cortical areas which locates both signals at successive stages in the cortical hierarchy. We show that cortical responses to sensory targets rely on activation patterns which are sustained in higher-order cortices across time only when sensory targets are complex and require the attentional tracking of the target for different sensory modalities, Conversely, the detection of targets which deviate from a short preceding stimulus stream and require only short-term memory produce a cortical signal which is propagated along cortical regions in a mid-latency time window regardless of sensory modality. Converging results from different temporal decoding analyses, we conclude that the prolonged maintenance of cortical activation elicited by the global ERP response and the serial propagation of the local ERP response are principles of cortical function found across sensory modalities. This demonstrates that cortical hierarchies implement target detection processes which track sensory irregularities in hierarchically nested different timescales at successive cortical processing stages for each sense.

### Rule-based sensory targets elicit supramodal and sustained responses in the cortex

Our finding that the sustained common supramodal activation patterns support the P3-like global ERP response contributes evidence to a controversy around its putative supramodal underpinnings. Some studies investigating cortical activation linked to the auditory and visual P3 suggest a common network including the insula and frontoparietal areas between these senses (Linden et al., 1999; Walz et al., 2013). However, other studies highlight a contribution of modality-specific higher-order regions to the P300 (Bledowski et al., 2004), leaving the question open whether and which sensory modalities share cortical networks to support P3-like global cortical signals. Our results support the notion that a supramodal (auditory, visual, somatosensory) network underpins the P3-like global ERP response while at the same time not ruling out contributions of modality-specific processes.

Complementing evidence that intrinsic neural timescales are linked to conscious information processing (Zilio et al., 2021), the global ERP response has also been proposed to be a cortical signal reflecting the conscious processing of incoming sensory stimulation (Bekinschtein et al., 2009). By this view, the global ERP response reflects recurrent information flow in a global neuronal workspace which maintains cortical signatures to become consciously accessible (Dehaene & Changeux, 2011). Sensory inputs from modality-specific cortices are fed forward to the global neuronal workspace which broadcasts integrated multisensory information from the top down to the levels below (Mashour, Roelfsema, Changeux, & Dehaene, 2020). Our observation that cortical signatures supporting the global ERP response are supramodal aligns with the theory that the global ERP response marks a supramodal top-down-driven process in which sensory information is amplified for conscious access via allocated attention (Chennu et al., 2013).

### Local ERP responses to sensory targets are linked to short-lived modality-specific activation

A classic view of MMN-like local ERP responses states that they rely on modality-specific cortical networks involving primary and secondary sensory regions (Nyman et al., 1990; Pazo-Alvarez et al., 2003). In our study, temporal cross-decoding analyses uncovered short-lived supramodal signatures for the local ERP response starting from ∼200 ms after onset of the last stimulus in a trial. Interestingly, previous studies demonstrate that local ERP responses are supported by a network involving modality-specific and frontal regions in which neuronal messages are propagated forward to the inferior frontal gyrus after initial processing in primary sensory areas, raising the possibility that frontal contributions to the MMN might host supramodal signatures. Indeed, both the visual and auditory MMN were found to consist of an earlier component in modality-specific early sensory cortices followed by an attention-modulated late frontal component from ∼200 ms after oddball onset (Deouell, 2007; Hedge et al., 2015). Similarly, studies of effective connectivity underpinning the MMN show that the potential is likely supported by a network spanning primary and secondary sensory cortices as well as frontal regions in different sensory modalities (Auksztulewicz & Friston, 2015; Chennu et al., 2016; Fardo et al., 2017; Garrido, Kilner, Kiebel, & Friston, 2009; Ostwald et al., 2012). Finally, the temporal characteristics of supramodal signatures supporting the local ERP response are congruent with a contribution of frontal areas linked to attention and target detection (Garrido et al., 2009a). Combined with earlier results, our results suggest that the MMN-like local ERP response might consist of short-lived modality-specific and supramodal components. In sum, this finding provides evidence for the idea that successive layers in the cortical hierarchy might support increasingly supramodal processes.

### A gradient of supramodality as a principle of cortical organisation

The canonical view of cortical function states that cortical hierarchies implement a strict unimodal-to-supramodal gradient. According to this view, supramodal processing is deferred to associative and frontal cortices (Felleman & Van Essen, 1991). Mounting evidence demonstrates that multisensory processes are ubiquitous in the cortical hierarchy and occur at all processing stages which refutes the idea that early cortices are strictly unimodal (Driver & Noesselt, 2008; Ghazanfar & Schroeder, 2006). Integrating both views, our findings support the view that the cortex is hierarchically organised along a gradient of supramodality. Earlier studies employed the local-global paradigm to demonstrate that the MMN-like local ERP response is generated in the primary auditory cortex whereas the global ERP response relies on activity in frontoparietal regions (Bekinschtein et al., 2009; Chao et al., 2018; Chennu et al., 2013; El Karoui et al., 2015; Uhrig, Dehaene, & Jarraya, 2014; Wacongne et al., 2011). In our study, feature-based sensory irregularities triggering quickly decaying and early modality-specific processes were supplemented by a supramodal contribution (Figure 6). Finally, a sustained late response to rule-based sensory irregularities shared between sensory modalities might reflect a recurrent supramodal process in higher-order cortical areas. However, our finding that local responses evolve into short-lived supramodal activation patterns provide evidence for the notion that early cortical function is not strictly specific to a sensory modality. Crucially, our finding that the P3b-like global ERP response relies on sustained supramodal cortical signatures while the local ERP response elicits early responses with short-lived commonalities between the senses supports the notion of a gradient of supramodality underpinning cortical hierarchies but also refutes the idea that early cortical target detection processes are strictly modality-specific.

**Figure 6.**
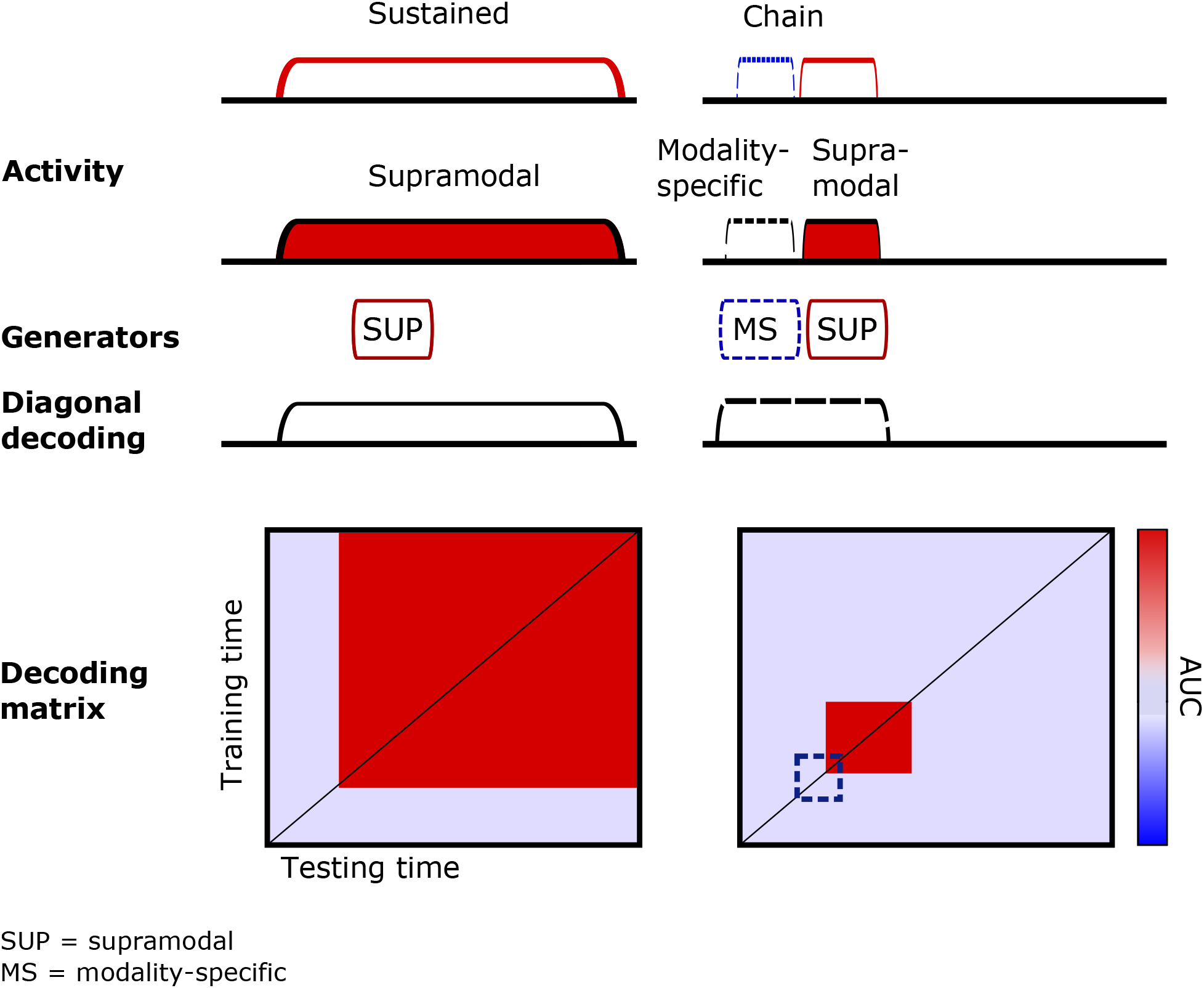
Summary. Supramodal and modality-specific aspects of the local and global ERP response. A temporal decoding analysis within sensory modality shows that the global ERP response is supported by one sustained process whereas the local ERP response likely relies on a chain of processes. Follow-up temporal decoding analyses from one modality to the other, and combining all sensory modalities, revealed that the global ERP response activates a single supramodal network across time whereas the local ERP response is propagated along the cortical hierarchy in a series of short-lived modality-specific and supramodal processes. We infer that a single supramodal generator contributes to the global ERP response whereas the local effect is likely supported by a chain of modality-specific and supramodal generators. Cortical activity indexing the global ERP response leads to an extended rectangular classification score matrix and some classification score clusters ordered along the diagonal. Supramodal processes found in temporal cross-decoding between sensory modalities are highlighted in red and modality-specific processes are shown in blue. We also show a list of supramodal and modality-specific properties of the local and global ERP response (King & Dehaene, 2014).

Finally, our results can be interpreted as evidence for a predictive coding view of cortical function. Predictive coding states that cortical responses to irregular sensory information reflect a prediction error resulting from a reconciliation of actual sensory inputs and their predictions (Clark, 2013; Friston, 2005; Hohwy, 2012; Rao & Ballard, 1999). From this perspective, local and global ERP responses can be seen as manifestations of prediction errors located at temporally dissociable successive levels of a dedicated cortical hierarchy for each sense (Wacongne et al., 2011). A central idea in predictive coding is that higher-order levels of the cortical hierarchy converge information from different senses forwarded from the levels below to generate predictions about the sensory environment (Clark, 2013; de Lange et al., 2018; Friston, 2005; Hohwy, 2012). This aligns with our result that higher-order cortical responses share supramodal signatures between the senses while lower-order responses largely rely on modality-specific activation patterns. Extending these earlier findings, we deliver an integrative framework for cortical responses to sensory targets tracking different time-scales at successive levels of the cortical hierarchy across different sensory modalities.

### Code accessibility

Code can be accessed here:

https://github.com/marianiedernhuber/Sustained-supramodal-signatures-target-detection

